# Genome-wide characterisation of the metallothionein gene family and their expression in response to copper in *Pistia stratiotes* (water lettuce)

**DOI:** 10.64898/2026.09.19.752864

**Authors:** Tapiwa Nyakauru, Orla O’Donovan, David O’Neill

## Abstract

Heavy metal contamination of aquatic environments poses persistent risks to ecosystems and human health. Aquatic macrophytes such as *Pistia stratiotes* are increasingly used for phytoremediation due to their capacity to accumulate contaminants without displaying symptoms of abiotic stress. However, the molecular mechanisms underlying their metal tolerance remain poorly understood. Metallothioneins (MTs) are cysteine-rich proteins involved in metal binding and detoxification in plants. In this study, genome-wide analysis was performed to identify and characterise candidate MT genes in *P. stratiotes* and to assess their transcriptional response to copper exposure. Six novel MT genes (*PsMT2f*, *PsMT3a-d*, and *PsMT4*) were identified and classified into MT2, MT3, and MT4 subfamilies based on conserved cysteine motifs and domain architecture. Quantitative real-time PCR showed differential and tissue-specific expression of MT genes in roots and leaves following exposure to copper for up to 4 h. These findings provide the first comprehensive characterisation of the MT gene family in *P. stratiotes* and demonstrate their potential role in copper tolerance, supporting the application of this species in the phytoremediation of metal-contaminated aquatic systems.

## 1 Introduction

Heavy metals occur naturally in the earth’s crust and cannot be broken down or destroyed. Their presence in water poses serious risks to humans, animals, and the environment (Morales *et al*., 2016; Tovar-Sánchez *et al*., 2018; Zhou *et al*., 2020). The increase in heavy metal contamination has been attributed to anthropogenic activities such as agriculture, mining, borehole drilling, construction, metal manufacturing, and leather production (Briffa *et al*., 2020). Furthermore, climate-related factors such as extreme weather events, increased temperatures, ocean acidification, and production of acid rain promote the leaching of heavy metals from rocks and soils, leading to higher levels of heavy metals in water systems (Li *et al*., 2024; Shu *et al*., 2024; Zheng *et al*., 2012).

Heavy metal contamination of water is commonly addressed through several traditional methods including electrochemical treatment, chemical precipitation, membrane filtration, ion exchange, and adsorption (Englande *et al*., 2015; Gunatilake, 2015; Padmavathiamma & Li, 2007; Wu *et al*., 2010). In recent years, plants have emerged as sustainable and cost-effective alternatives for heavy metal remediation, owing to their natural capacity to uptake, translocate, and stabilise metal ions from contaminated sites (Kumar *et al*., 2024). Among these, aquatic plants have attracted significant attention due to their continuous interaction with the water, which facilitates phytoaccumulation (Lu *et al*., 2018). Hyperaccumulator aquatic plants possess a remarkable capability for heavy metal adsorption and absorption without showing signs of abiotic stress (Kumar *et al*., 2018; Ugya *et al*., 2015). Of particular importance to this study is *Pistia stratiotes* (water lettuce), a free-floating aquatic macrophyte native to tropical and subtropical regions, which originated from South America. It thrives in slow-moving or stagnant freshwater habitats, including ponds, lakes, and rivers. While the rapid growth of *P. stratiotes* may pose significant ecological risks, it facilitates the formation of dense biomass that enhances its capacity for the uptake, accumulation, and tolerance of contaminants from aquatic environments (Kurugundla *et al*., 2016; Pang *et al*., 2023; Šajna *et al*., 2007).

The phytoremediation potential of *P. stratiotes* has been widely demonstrated across laboratory and pilot-scale studies. When exposed to three concentrations (1.0, 2.0, and 5.0 mg/L) of five heavy metals (Fe, Zn, Cu, Cr, and Cd), *P. stratiotes* accumulated more than 90% of the metals present within 15 days (Mishra & Tripathi, 2008). In contaminated waters and industrial effluents, at metal concentrations ranging from 5 to 20 mg/L, *P. stratiotes* achieved 80-93% removal of Pb and Cr within 10-20 days (Abubacker & Sathya, 2016; Zhou *et al*., 2013). Furthermore, exposure to 10 mg/L of Pb, Zn, Cr, Cu, and Ni for 25 days resulted in overall removal efficiencies of 73-84%, with Pb exhibiting the highest uptake (Samal & Dash, 2024). Recent studies are performing pilot experiments of using *P. stratiotes* in phytoremediation. In a 40-day phytoremediation study using glass industry effluent containing 1-5 mg/L of Cu, Cr, Fe, Mn, Pb, and Zn, *P. stratiotes* showed optimal performance at 25% effluent dilution, achieving high metal removal efficiencies of Cr (95.29%), Cu (91.74%), Zn (91.34%), Mn (92.95%), Pb (87.10%), and Fe (86.47%), demonstrating its strong capacity for heavy-metal accumulation and remediation of industrial wastewater (Singh *et al*., 2024). Beyond contaminant removal, the substantial accumulation of metals within *P. stratiotes* biomass presents opportunities for resource recovery. Rather than treating the harvested biomass solely as waste, the accumulated metals may be recovered through biomining or related metal recovery approaches, supporting the transition from conventional wastewater treatment towards more circular and sustainable resource management (Richardson & Mirkouei, 2026).

Heavy metal uptake by plants involves root adsorption, absorption, translocation, and sequestration, which largely depends on various genes, proteins, and regulatory components that are known to be activated in the presence of heavy metals (Rai *et al*., 2020; Yan *et al*., 2020). Plants rely on intracellular detoxification mechanisms that restrict the interaction of free metal ions with essential cellular components. They employ several classes of metal-binding molecules for detoxification, including glutathione, phytochelatins, and metallothioneins. Metallothioneins (MTs) are of particular interest because they are directly encoded by genes and are known to be strongly induced in response to heavy metal exposure, making them key components of metal homeostasis and tolerance pathways. They typically have a low molecular weight from 4-8 kDa and contain highly conserved cysteine-rich motifs that contribute close to 30% of the polypeptide chain (Binz & Kägi, 1999; Leszczyszyn *et al*., 2013). The abundance of cysteine residues provides sulfhydryl (-SH) groups that enable MTs to bind heavy metal ions, forming stable protein-metal complexes that can subsequently be transported into vacuoles for storage (Gu *et al*., 2020; Joshi *et al*., 2016; Zhao *et al*., 2022).

The capacity of *P. stratiotes* to hyperaccumulate heavy metals suggests the potential involvement of efficient intracellular detoxification and sequestration mechanisms of heavy metals in *P. stratiotes*. However, the molecular mechanisms of these processes in *P. stratiotes* during heavy metal accumulation are poorly understood, largely due to the limited availability of nucleotide sequence information. In this study, we employed genome wide analysis to screen for novel MTs in *P. stratiotes* chromosomes and we identified six novel MTs and three endogenous control genes. Relative gene expression analyses showed a varying expression pattern of novel MTs, collectively confirming their involvement in heavy metal tolerance and accumulation in *P. stratiotes*. This tissue-specific response pattern highlights that different MTs might have distinct roles in detoxifying heavy metals in different plant parts. Our results provide crucial information on the evolution of the MT gene family in *P. stratiotes* and improves our understanding of the role of *P. stratiotes* MTs gene in conferring copper stress tolerance.

## 2 Methodology

### 2.1 Identification of candidate MT genes

BLASTp analyses against local MT databases were performed using *P. stratiotes* polypeptide sequences deposited at the China National GeneBank Database by Chen *et al*., (2020). Sequences with similarities to published MTs were noted, and their coding sequences were identified in the *P. stratiotes* mRNA sequences deposited under accession CNS0457649 (Qian *et al*., 2022).

To identify unannotated MT genes, six-frame translation was performed on the *P. stratiotes* chromosomes using Geneious Prime 2023.1.2. Conserved MT domains were manually screened within the translated polypeptides. Conserved cysteine-rich motifs characteristic of MT1-MT4 proteins were defined as follows: MT1, CxCxxxCxCxxxCxC, CxCxxxCxCxxCxC; MT2: CCxxxCxCxxxCxCxxxCxxC, CxCxxxCxCxxCxC, KYMPD, and MSCCG; MT3: CxxCxCxxxxxC, CxCxxxCxCxxCxC, MADTGKS; and MT4: CxxxCxCxxxCxxxxxCxC, CxCxxxCxCxxCxC, CxCxxxCxCxxC (where x represents any amino acid). Sequences containing MT conserved domains were retrieved for subsequent bioinformatic analyses.

### 2.2 Nucleotide sequence analysis

Gene mapping was performed using novel sequences as query sequences against *P. stratiotes* chromosomes for the identification of the complete genes, transcription start site, and gene loci, which was visualized using a mapchart (Voorrips, 2002) and Gene Structure Display Server (Hu *et al*., 2015). The presence of Kozak sequences was assessed (Kozak, 1987; and Kim *et al*. 2014), and the ratio of Ka/Ks was calculated using DNASP6 (Rozas *et al*., 2017) to predict duplication events of novel genes.

### 2.3 Polypeptide sequence analysis

BLASTp analyses were performed against the non-redundant protein database on NCBI (Altschul *et al*., 1990). Novel sequences were named as suggested by Cobbett & Goldsbrough (2002) and were deposited to GenBase (Bu *et al*., 2024) of the National Genomics Data Center (Bao *et al*., 2025). Conserved domains were accessed using the NCBI conserved domain database (NCBI-CDD). Multiple sequence alignment was performed on MEGA-X using published MT sequences from various plants and alignments were visualized on Jalview (Waterhouse *et al*., 2009). The physical and chemical properties were predicted using the ProtParam tool (Gasteiger *et al*., 2005) and subcellular localization was predicted using DeepLoc-1.0 (Goldberg *et al*., 2014). Multiple Expectation Maximization for Motif Elucidation (MEME tool) was used to identify conserved motifs in polypeptide sequences using the following parameters: number of repetitions, any; maximum number of motifs, 15; and optimum width of each motif between six and 300 residues (Bailey *et al*., 2009; Pan *et al*., 2018). MEGA-X was used to construct a Maximum Likelihood method and JTT matrix-based model and bootstrap method with 1,000 repeats to illustrate evolutionary relationships. The phylogenetic tree was rooted using the *Rhodobacteraceae* MT sequence (accession: X6L243) as an outgroup to provide evolutionary distance (Niu *et al*., 2018; Pan *et al*., 2018; Yang *et al*., 2015).

### 2.4 PCR primer design and optimisation

Real-time PCR primers were designed for *P. stratiotes* using the NCBI primer BLAST tool (Ye *et al*., 2012) to target the putative novel MT genes. Primers were also designed for endogenous control genes. The *18S rRNA* gene (accession: AH001726) was obtained from previous data (Davies *et al*., 2004), while β-actin, *GAPDH*, and α*-tubulin* were identified and characterised in this study from sequences deposited by Qian *et al*. (2022) using the methodology described in section 2.1. The primers were optimised for high specificity and efficiency to the target sequences, and had a GC content of 40-60%, annealing temperature of 60 °C, length of 18-24 base pairs, and amplicon size of 90-125 bp. Primers were analysed for secondary structures including hairpins, self-dimers, and cross-dimers using Eurofins genomics oligo analysis tool (https://eurofinsgenomics.eu/). The qPCR efficiency, specificity, and sensitivity of the novel primers were assessed based on standard curves, melt curves, and amplification curves, as described by Svec *et al*. (2015). Real-time PCR was performed using a QuantStudio 3 Real-Time PCR System (Applied Biosystems). PCR amplification targeting endogenous genes and MTs was conducted using 10-fold cDNA template dilutions between 10^0^-10^-6^ to achieve a minimum of four to five amplification curves. Each PCR reaction contained 5 μL 2X FastStart Universal SYBR® Green Master Mix (ROX) (Catalogue number: 04913914001), 100 nM forward primer, 100 nM reverse primer, 1 μL template, and nuclease-free water to a final volume of 10 μL. All samples and controls were assayed in triplicate. The qPCR reaction protocol included 50 °C for 2 minutes and 95 °C for 10 minutes, followed by 40 cycles of 95 °C for 15 s and 60 °C for 1 minute. Following PCR, a melting curve was recorded by increasing the temperature at 0.15 °C/s from 60 °C to 95 °C. Primers with an acceptable efficiency of 100 ± 10% were selected for gene expression analysis.

### 2.5 Gene expression analysis by quantitative real time PCR

*P. stratiotes* plants were grown in the laboratory in 10% Hoagland solution (1 mM KH_2_PO_4_, 5 mM KNO_3_, 5 mM Ca[NO_3_]_2_, 2 mM MgSO_4_, 10 µM Fe-EDTA, 50 µM H_3_BO_3_, 9 µM MnCl_2_·4H_2_O, 0.77 µM ZnSO_4_·7H_2_O, 0.32 µM CuSO_4_·5H_2_O, and 0.12 µM H_2_MoO_4_·H_2_O) (Hoagland & Arnon, 1950; Peng *et al*., 2024; Singh *et al*., 2019) at 22 °C under a 16 h photoperiod supplied by 6700 K fluorescent lights with a light intensity of approximately 200 μmol m ² s ¹. The Hoagland solution was changed every seven days to replenish nutrients and maintain a constant pH, and any plant debris was removed at the same time to maintain clean growing conditions.

Plants of similar age with an average mass of 6.2 g were acclimatised for 14 days in 10% Hoagland solution. Following that, plants were exposed to 5 mg/L Cu, and sampling was performed at 0, 1, and 4 h. Each time point consisted of three independent biological replicates (n = 3). Control plants (n = 3) that were not exposed to Cu were included and processed under the same conditions. Following Cu exposure, plants from each biological replicate were harvested separately, and roots and leaves were separated and homogenised independently. The homogenised root and leaf samples from each biological replicate were immediately frozen in liquid nitrogen and stored at -80 °C. Three independent RNA extractions were performed from each homogenised biological sample as technical replicates using the RNeasy Plant Mini Kit (Qiagen, Catalogue number: 74904) and was further purified using the DNA max kit (Qiagen, Catalogue number: 15200-50). In total, 90 RNA extractions were performed, and the purity and concentration of total RNA was measured using a Nanodrop ND-8000 spectrophotometer. RNA concentrations across all samples were standardised for reverse transcription, and first-strand cDNAs were synthesised from total RNA using a High-Capacity RNA-to-cDNA Kit (Thermofisher, catalogue number: 4388950). Quantitative PCR was conducted as mentioned before, using validated primer pairs on cDNA synthesised from control and treated RNA samples.

### 2.6 Reference gene expression stability

Threshold cycle (Ct) values for all candidate reference genes were used to construct box and whisker plots (Burns *et al*., 2005). The mean, standard deviation (SD), and coefficient of variation (CV) were calculated for each of the three biological replicates, as described by Zhao *et al*. (2022). Gene expression stability was also calculated using BestKeeper (Köhsler *et al*., 2020; Pfaffl *et al*., 2004), NormFinder (Andersen *et al*., 2004), and Genorm (Vandesompele *et al*., 2002) statistical tools integrated into the RefFinder web-based tool (Xie *et al*., 2023).

### 2.7 Relative quantification of target genes

The fold change in the expression of target genes was calculated from the Ct values of the target and reference genes in treated and control samples using the comparative 2^−ΔΔ*Ct*^ method (Livak & Schmittgen, 2001), where ΔΔCt = ΔCt_treated_ -ΔCt_calibrator_, ΔCt_treated_ = Ct_target_ -Ct_normalizer_, and ΔCt_calibrator_ = Ct _target_ -Ct_normalizer_. The most stable candidate reference gene was used in relative gene expression calculations (Andersen *et al*., 2004; Pfaffl *et al*., 2004; Vandesompele *et al*., 2002). GraphPad Prism was used for statistical analysis, and the significance of the results was calculated using two-way ANOVA (p<0.05) (Mousavi *et al*., 2021).

## 3 Results

### 3.1 MTs sequence identification sequence analyses

In this study, we successfully identified six candidate MT genes containing conserved metallothio_2 and metallothio_PEC domains (Pan *et al*., 2018), and these were named *PsMT2f*, *PsMT3a-d*, and *PsMT4,* following the nomenclature convention proposed by Binz & Kägi (1999) and Cobbett & Goldsbrough (2002). Amino acid percentage identity ranging from 53% to 88% was observed when comparing these sequences with previously published MT sequences (**Table 3.1**).

**Table 3.1:** Characteristics of novel MT Genes Identified in *P. stratiotes* and their sequence-related features.

|  |  |  |  |  |  |  |  |  |  |  |  |  |  |  |  |  |
| --- | --- | --- | --- | --- | --- | --- | --- | --- | --- | --- | --- | --- | --- | --- | --- | --- |
| Sequence | Accession | Accession | Gene | Exons | Introns | <u>Kozak</u> |  | CDS | Length | MW | pI | Hydropathy | Subcellular | Best BLASTp hit | Identity (%) | Accession |
| name | (Nucleotide) | (protein) | (bp) |  |  | -3(R) | +4(G) | (bp) | (aa) | (kDa) |  | index | localization |  |  |  |
| ***PsMT2f*** | C_AA103793.1 | C_AAK65816.1 | 957 | 3 | 2 | A | T | 246 | 81 | 8.03 | 4.49 | 0.08 | Cytoplasm | *Colocasia esculenta* [MT2B] | 88.33% | Q19LA2.1 |
| ***PsMT3a*** | C_AA103794.1 | C_AAK65817.1 | 624 | 3 | 2 | A | T | 195 | 64 | 6.61 | 5.44 | -0.3 | Extracellular | *Typha angustifolia* [MT3] | 67.19% | ACV51811.1 |
| ***PsMT3b*** | C_AA103795.1 | C_AAK65818.1 | 666 | 3 | 2 | G | T | 198 | 65 | 6.77 | 5.08 | -0.36 | Extracellular | *Colocasia esculenta* [MT3] | 78.46% | MQM09729.1 |
| ***PsMT3c*** | C_AA103796.1 | C_AAK65819.1 | 717 | 3 | 2 | G | T | 198 | 65 | 6.77 | 5.08 | -0.36 | Extracellular | *Colocasia esculenta* [MT3] | 78.46% | MQM09729.1 |
| ***PsMT3d*** | C_AA103797.1 | C_AAK65820.1 | 717 | 3 | 2 | G | T | 198 | 65 | 6.79 | 5.08 | -0.41 | Extracellular | *Colocasia esculenta* [MT3] | 78.46% | MQM09729.1 |
| ***PsMT4*** | C_AA103798.1 | C_AAK65821.1 | 514 | 2 | 1 | G | G | 294 | 97 | 9.36 | 6.94 | -0.39 | Extracellular | *Spirodela intermedia* [MT4] | 53.61% | CAA7410690.1 |

Alignment of candidate MTs with sequences from model plants such as *Arabidopsis thaliana*, *Zea mays*, and *Oryza sativa* show conserved cysteine arrangements typical of plant MTs. *PsMT2f*, consisting of 81 amino acids had cysteine motifs of CCxxxCxCxxxCxCxxxCxxC in the N-terminal region and CxCxxxCxCxxCxC in the C-terminal end, and was classified as type 2. *PsMT3a-d* consisted of 64-65 amino acids, had homology ranging between 58-100%, and had their cysteine motifs arranged as CxxCxCxxxxxC in the N-terminal and CxCxxxCxCxxCxC in the C-terminal, typical of type 3 MTs. *PsMT4* had 97 amino acids and its cysteine residues arranged in three distinct motifs of CxxxCxCxxxCxxxxxCxC, CxCxxxCxCxxCxC, and CxCxxxCxCxxC, and was classified as type 4 (**Figure 3.1**). Novel candidate MTs were deposited in the National Genomics Data Center (https://ngdc.cncb.ac.cn/genbase/) under accession C_AA103793.1-C_AA103798.1.

**Figure 3.1:**
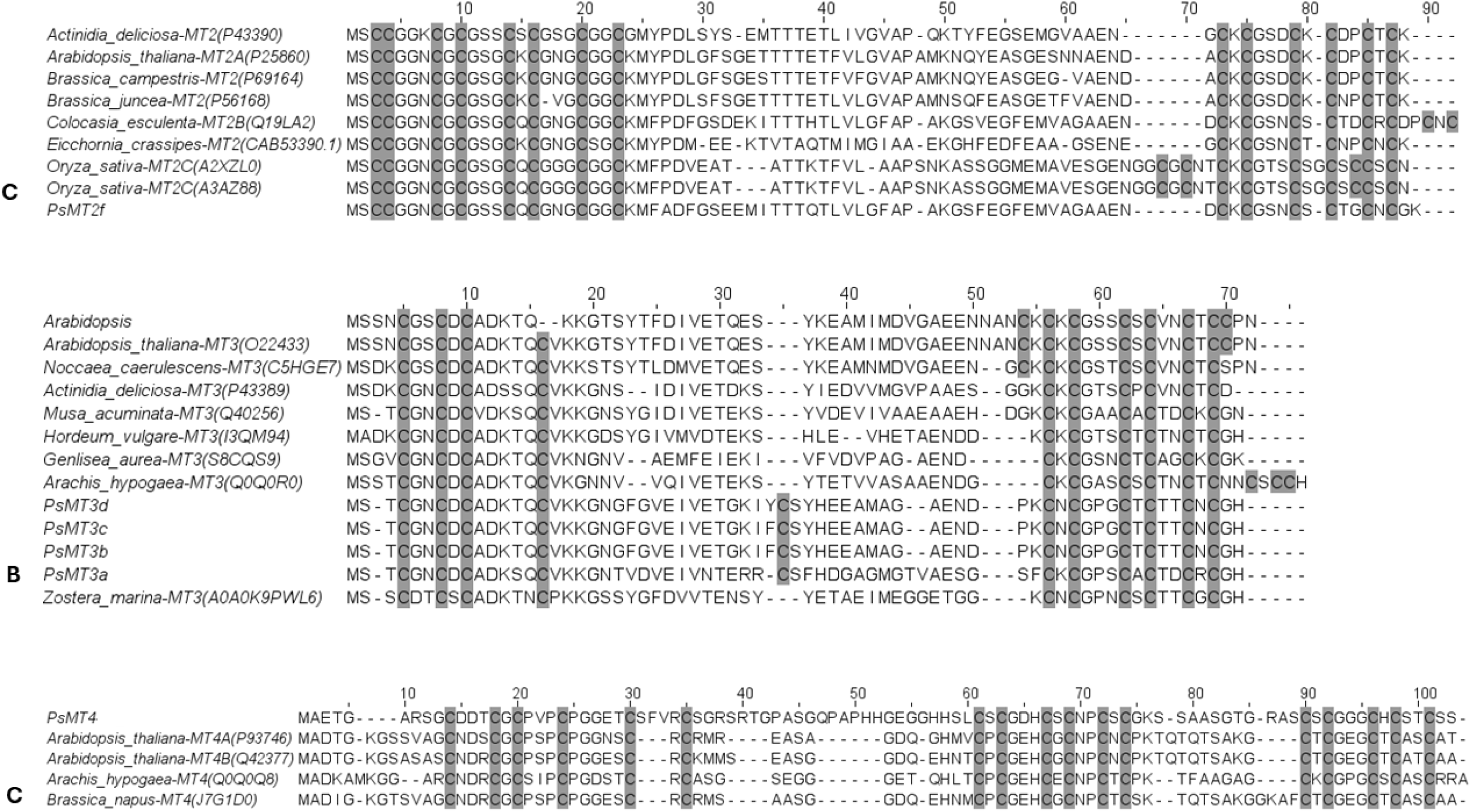
Multiple sequence alignment of novel putative (A) MT2, (B) MT3, and (C) MT4 sequences from *P. stratiotes* with published sequences from other plant species.

Amino acid residues that form the linker region that connects metallothio_2 and metallothio_PEC domains varied, with some linker regions containing aromatic amino acids such as tyrosine, histidine, and phenylalanine (**Figure 3.1**). Isoelectric point (pI) of novel MTs were in the range of 4.49-6.94, and hydropathy index in the range of -0.41-0.08 (**Table 3.1**). Using DeepLoc, *PsMT2f* was predicted to be located in cytoplasm while *PsMT3a-d* and *PsMT4* were predicted to be extracellular proteins.

A neighbour-joining phylogenetic tree revealed that *PsMT2f* clustered closely with previously published MT2 sequences while *PsMT3a-d* and *PsMT4* formed a distinct cluster with MT3 and MT4 sequence, respectively, from plants such as *A. thaliana, Hordeum vulgare,* and *Arachis hypogaea* (**Figure 3.2**).

**Figure 3.2:**
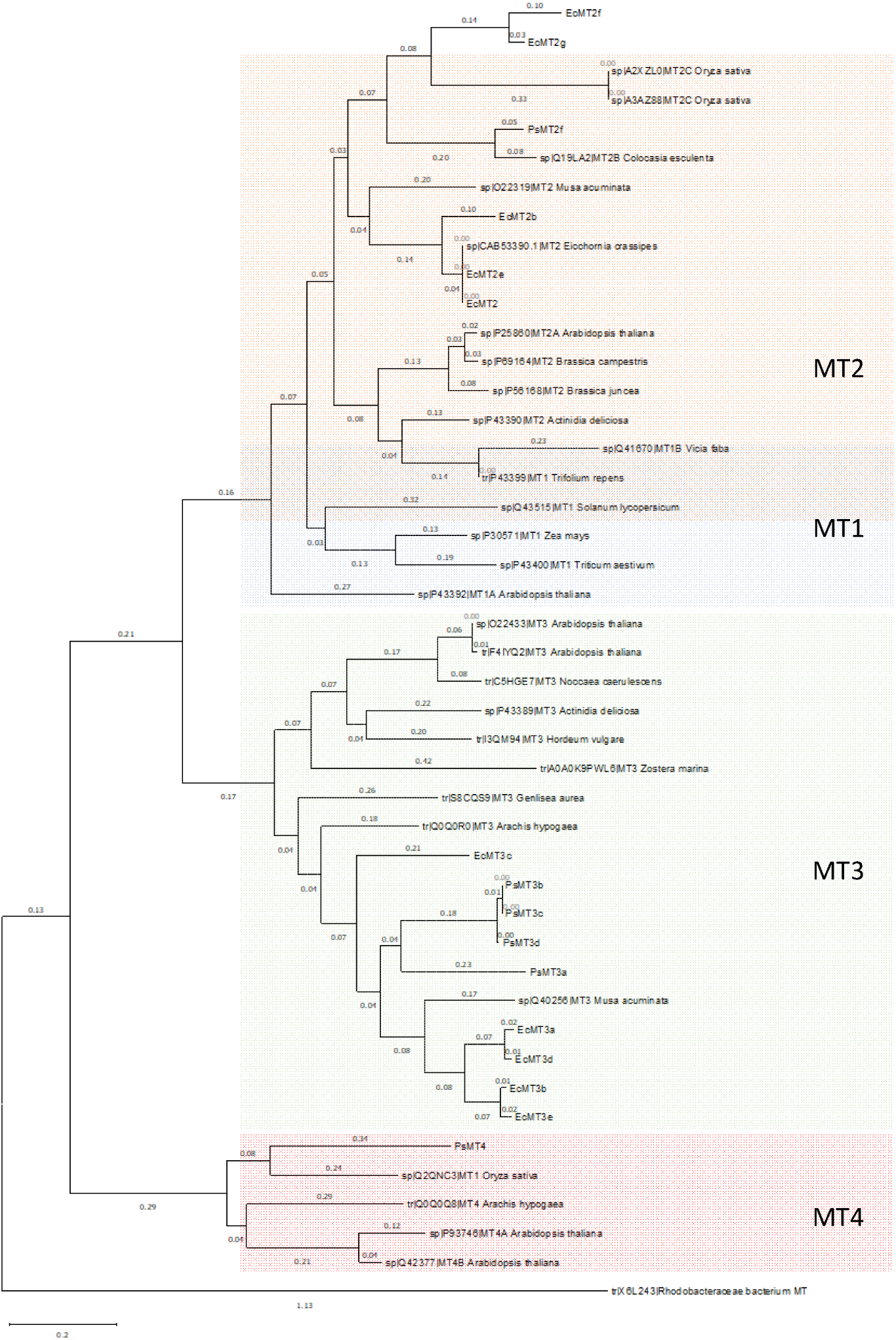
A phylogenetic tree showing the evolutionary relationships between novel putative MTs and published MT sequences from various plant species constructed using MEGA X. The MT sequence from Rhodobacteraceae was used as the outlier.

### 3.2 Genomic structure and gene duplication

The gene structures of the MT family genes were characterised by comparing the CDS and the genomic DNA sequences published by Qian *et al*. (2022) using Gene Structure Display Server (Hu *et al*., 2015). *PsMT2f* had three exons, *PsMT3a-d* had three exons, and *PsMT4* had two exons (**Figure 3.3**). The analyses showed a high proportion of predicted intronic regions in novel MTs (42-74%). Analysis of Kozak sequences showed that *PsMT2f* and *PsMT3a-d* contained a –3 (R) nucleotide and a +4 (T) nucleotide, whereas PsMT4 had a -3R nucleotide and a +4G nucleotide from the transcription start site (**Table 3.1**).

**Figure 3.3:**
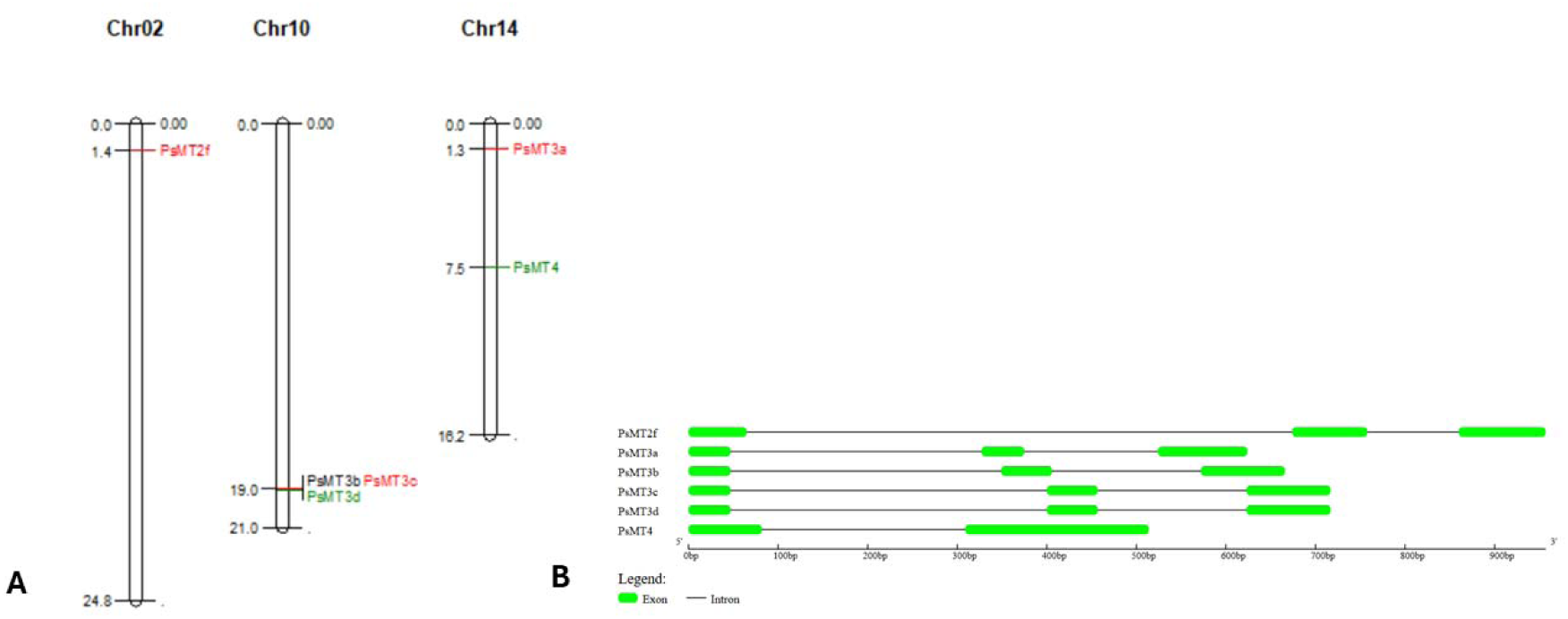
(A) Chromosomal distribution and analysis of novel putative MT genes in *P. stratiotes*. The lengths of the chromosomes are in megabases (Mb); (B) Genomic structures of the novel MTs. The green boxes represent exons, and the solid black lines represent introns.

To investigate gene duplication events in *P. stratiotes*, the ratio of Ka (the nonsynonymous substitution rate) to Ks (the synonymous substitution rate) was calculated and results are presented in **Table 3.2**.

**Table 3.2:** Ka/Ks analysis of novel MTs.

| Gene | Sequence 1 | Sequence 2 | Ka | Ks | Ka/Ks |
| --- | --- | --- | --- | --- | --- |
| *P. stratiotes* MT3 | *PsMT3a* | *PsMT3b* | 0.25 | 0.41 | 0.60 |
|  | *PsMT3a* | *PsMT3c* | 0.25 | 0.41 | 0.60 |
|  | *PsMT3a* | *PsMT3d* | 0.25 | 0.41 | 0.60 |
|  | *PsMT3b* | *PsMT3c* | 0 | 0 | 0 |
|  | *PsMT3b* | *PsMT3d* | 0 | 0 | 0 |
|  | *PsMT3c* | *PsMT3d* | 0 | 0 | 0 |

### 3.3 Motif and cis-regulatory elements analyses

Using the MEME online tool, analysis of the MT3 sequences revealed the presence of three conserved motifs 1, 2, and 3, indicating a consistent structural pattern among these sequences. In contrast, *PsMT2f* and *PsMT4* displayed a greater diversity of motifs, including motifs 4-15 as presented in **Figure 3.4**.

**Figure 3.4:**
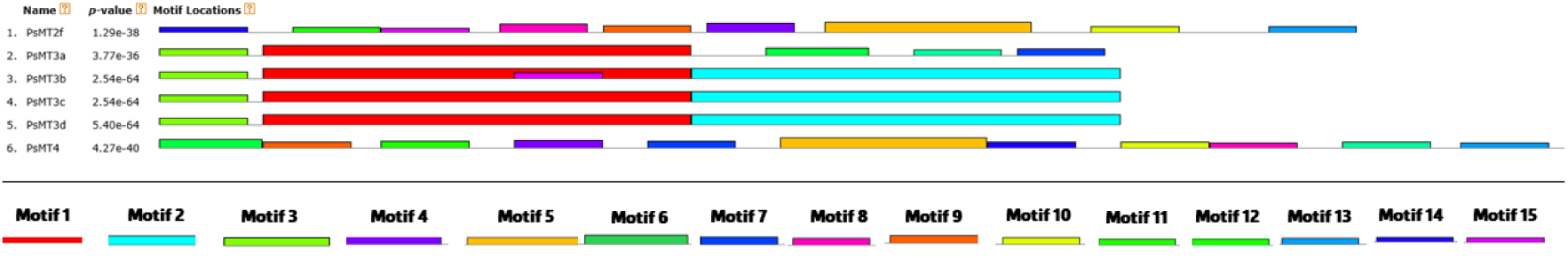
Putative conserved motifs in MT family proteins from *P. stratiotes* identified using the MEME search tool.

### 3.4 Assessment of primer specificity and amplification efficiency **Table 3.3**: Details of primers used for real time PCR

**Table 3.3:**
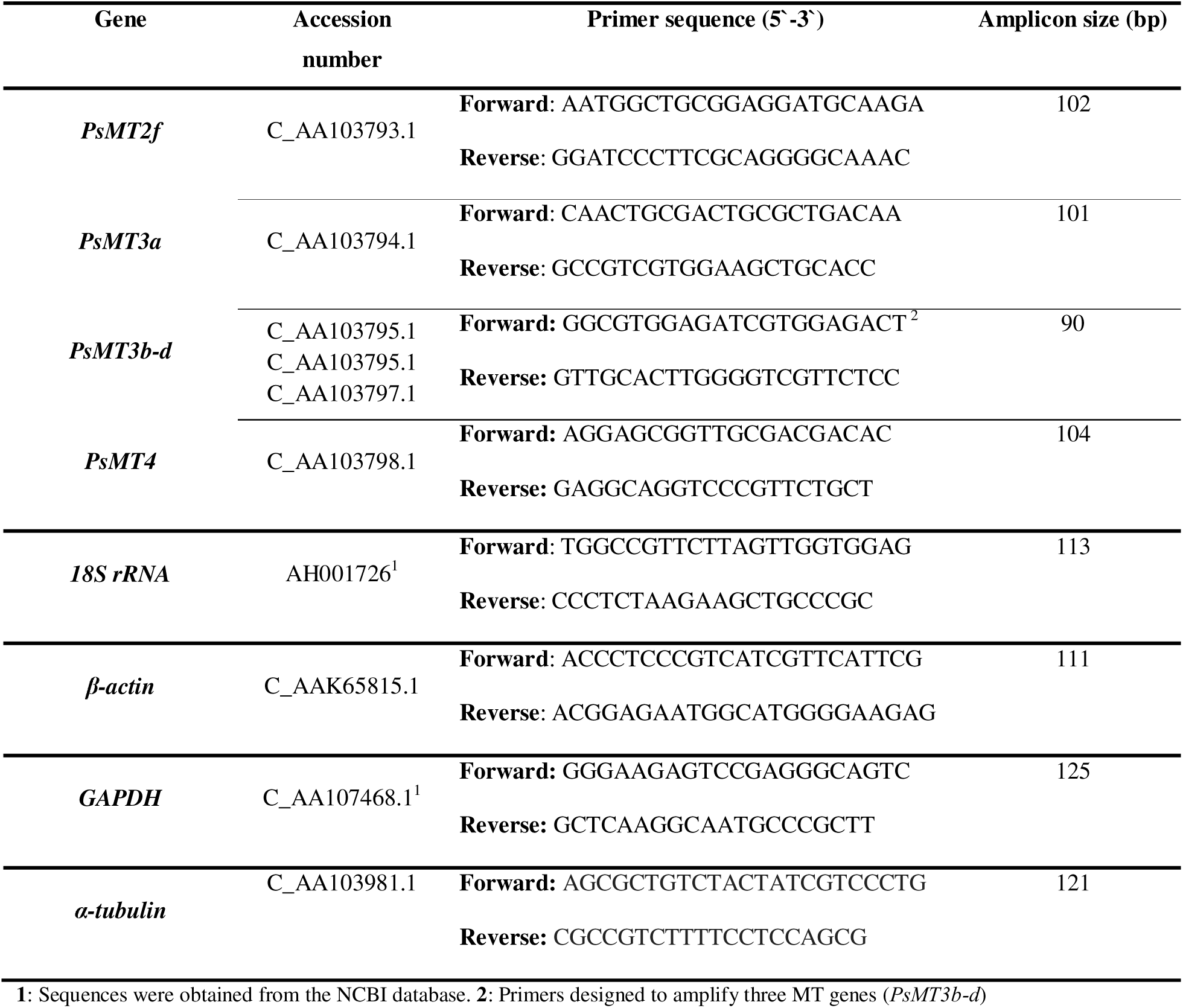
Details of primers used for real time PCR.

Primers designed to amplify *PsMT2f*, *PsMT3b-d*, *18S rRNA*, and β*-actin* successfully amplified the target genes as illustrated by the amplification curve and melt curve in **Figure 3.5**, and had expected primer efficiency in the range of 100 ± 10% (**Table 3.4**) (Ginzinger, 2002; Livak & Schmittgen, 2001). Melt curve analysis showed single peaks, and agarose gel showed single amplicons of expected size. The primers for α*-tubulin*, *GAPDH*, *PsMT3a*, and *PsMT4* showed less than the required number of amplification curves, suggesting that while amplification occurred, target genes were not expressed sufficiently to allow for primer optimization and calculation of primer efficiencies (Lorenz, 2012), therefore, they were not used in further experiments.

**Table 3.4:**
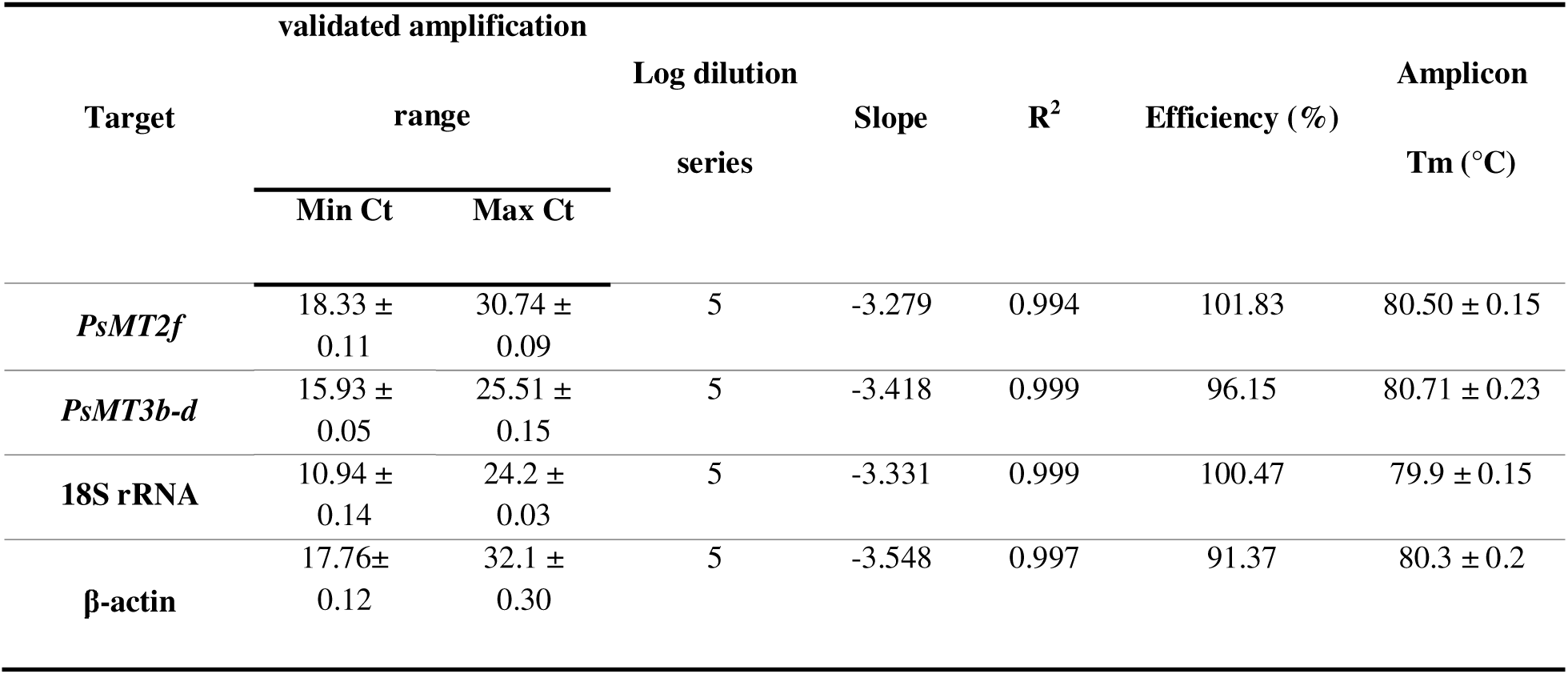
Quantitative real time PCR using novel primers to amplify target genes in *P. stratiotes*.

| validated amplification |  |  | Log dilution series | Slope | R<sup>2</sup> | Efficiency (%) | Amplicon Tm (°C) |
| --- | --- | --- | --- | --- | --- | --- | --- |
| Target | range |  |  |  |  |  |  |
|  | Min Ct | Max Ct |  |  |  |  |  |
| *PsMT2f* | 18.33 ± 0.11 | 30.74 ± 0.09 | 5 | -3.279 | 0.994 | 101.83 | 80.50 ± 0.15 |
| *PsMT3b-d* | 15.93 ± 0.05 | 25.51 ± 0.15 | 5 | -3.418 | 0.999 | 96.15 | 80.71 ± 0.23 |
| 18S rRNA | 10.94 ± 0.14 | 24.2 ± 0.03 | 5 | -3.331 | 0.999 | 100.47 | 79.9 ± 0.15 |
| β-actin | 17.76± 0.12 | 32.1 ± 0.30 | 5 | -3.548 | 0.997 | 91.37 | 80.3 ± 0.2 |

**Figure 3.5:**
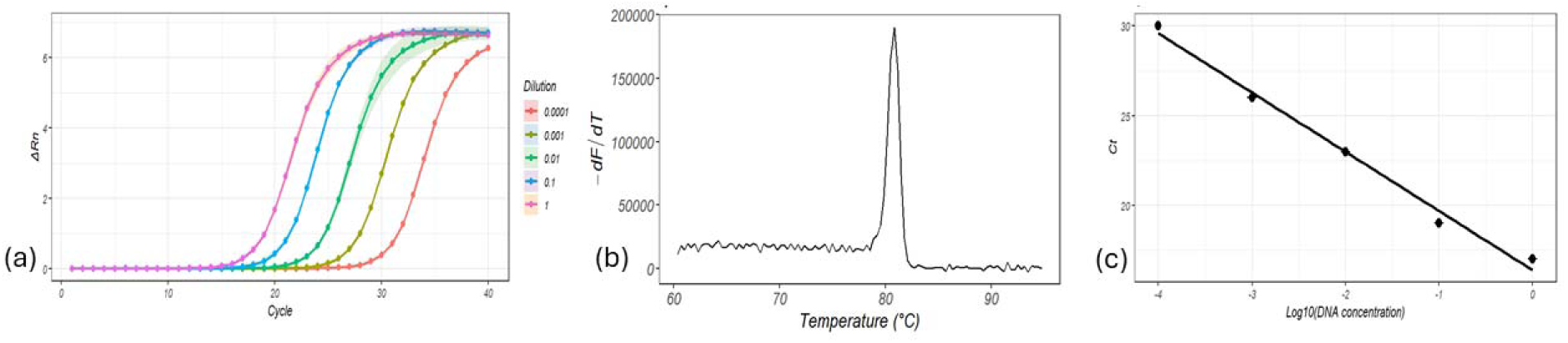
(a) Amplification plot showing a 1:10 serial dilution series of *P. stratiotes* cDNA with corresponding amplification curves, (b) melt curve plot showing a single peak of expected Tm, and (c) standard curve showing the linear regression of Ct values against DNA concentration, demonstrating assay efficiency and sensitivity

### 3.5 Normalization and statistical analysis of real time PCR results

To identify the most suitable reference gene for *P. stratiotes*, the expression stability of *18S rRNA* and β*-actin* was analysed in control plants and in plants exposed to 5 mg/L Cu. Box and whisker plots were constructed using Ct values (**Figure 3.6**) and the average Ct values for 18S rRNA were 9.83 ± 3.28 in leaves and 11.27 ± 4.37 in roots, while β*-actin* had average Ct values of 24.38 ± 2.27 in leaves and 23.92 ± 2.70 in roots.

**Figure 3.6:**
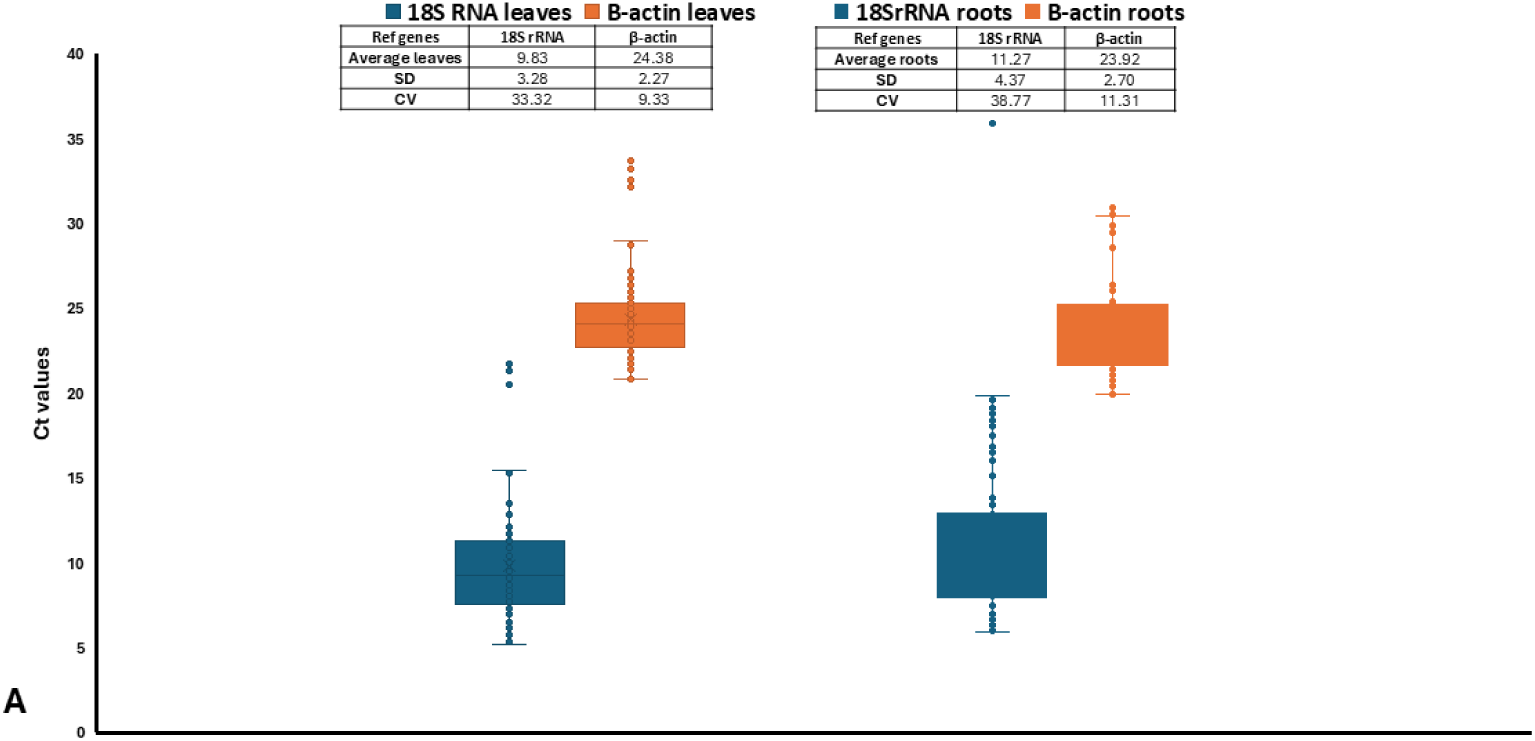
Ct values of reference genes 18S rRNA primers and β*-actin* in *P. stratiotes* exposed to Cu for 0-4h.

Further analysis of the stability of *18S rRNA* and β*-actin* gene expression was performed using the RefFinder web-based tool which combine BestKeeper, NormFinder, and Genorm tools, designed for expression stability analysis of reference genes. In our analysis using geNorm (**Table 3.5**), the obtained M value was >1.5 with both genes, and BestKeeper showed an SD of >1. Using NormFinder, the stability value obtained between β*-actin* and *18S rRNA* was 1.02 and 1.86 in roots and leaves, respectively, and the comprehensive ranking method, which combines the results from all three statistical tools showed that β*-actin* was the most stable gene in *P. stratiotes*.

**Table 3.5:**
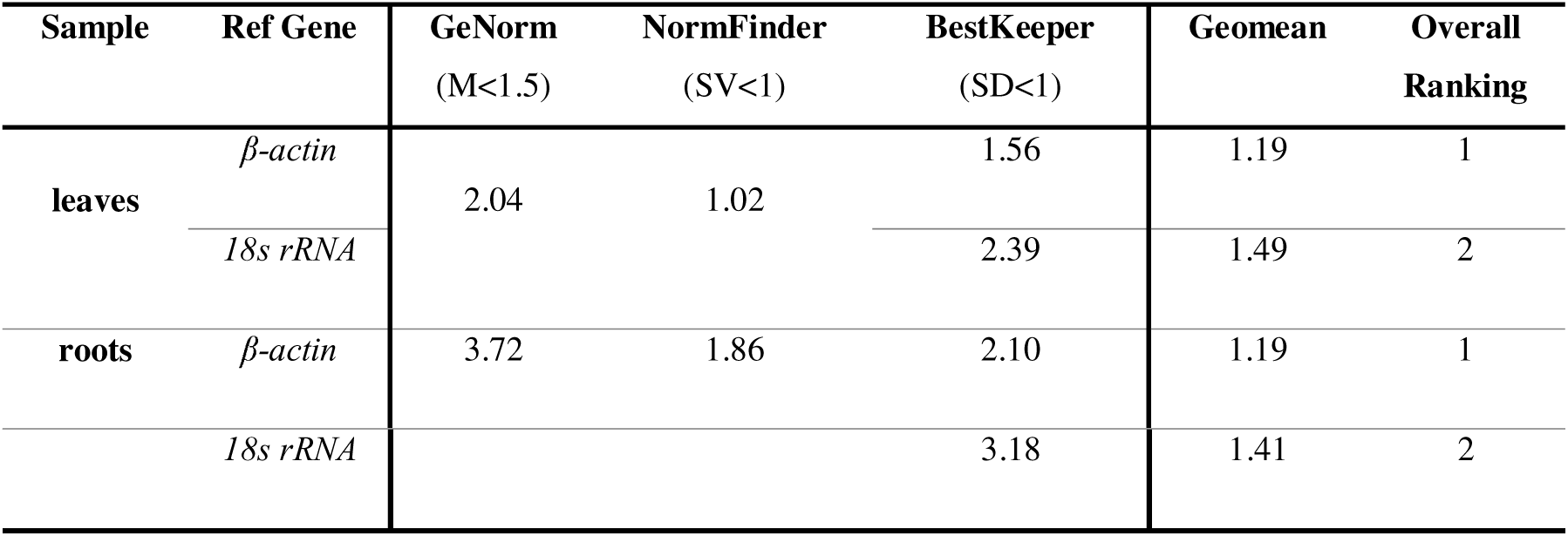
Gene expression stability of candidate reference genes ranked by geNorm, NormFinder, and BestKeeper, after exposure of *P. stratiotes* to 5 mg/L Cu for up to 4 h.

### 3.6 Expression profile of MTs under heavy metal stress

At 0 h, *PsMT2f* and *PsMT3b-d* were shown to be constitutively expressed in *P. stratiotes*. Notably, *PsMT2f* was expressed approximately 14-fold higher in leaves and 1.34-fold higher in roots than *PsMT3b-d*. At the 1 h time point in roots, *PsMT2f* expression slightly decreased to 0.87-fold, while *PsMT3b–d* showed a 1.71-fold increase. However, neither change was statistically significant (*p* > 0.05). In leaves, *PsMT2f* expression remained relatively unchanged at 1.32-fold (*p* > 0.05), whereas *PsMT3b–d* exhibited a significant upregulation, increasing to 3.21-fold (*p* < 0.05) (**Figure 3.7**). At the 4 h time point in roots, *PsMT2f* expression was significantly downregulated to 0.31-fold (*p* < 0.05), as shown in **Figure 3.7**, while *PsMT3b–d* expression showed a non-significant change at 0.69-fold (*p* > 0.05). In leaves, *PsMT2f* was significantly upregulated to 1.56-fold (*p* < 0.05), whereas *PsMT3b–d* exhibited a slight, non-significant change at 0.94-fold (*p* > 0.05) (**Figure 3.7**).

**Figure 3.7:**
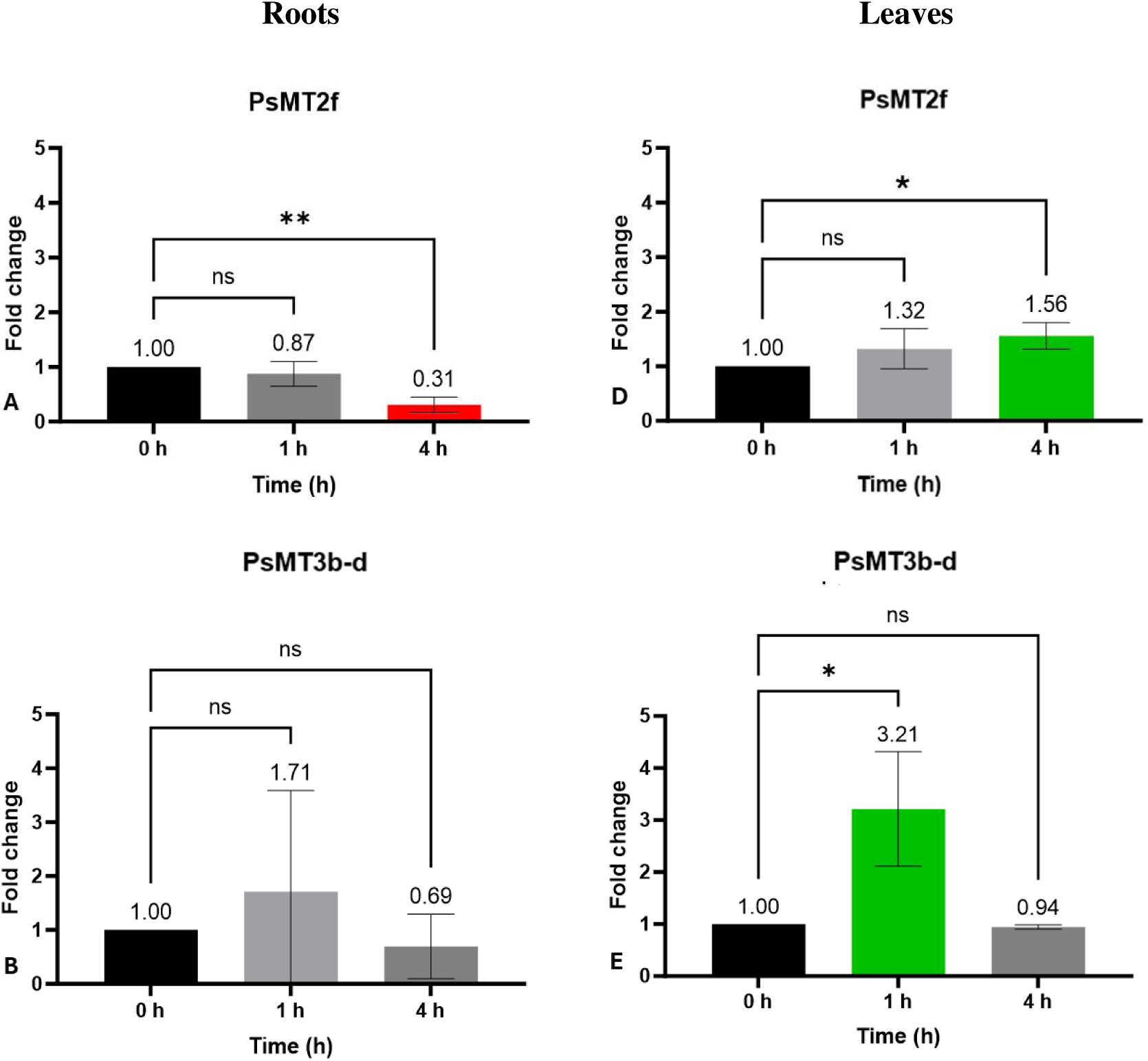
Relative expression levels of *PsMT2f* (A) and *PsMT3b-d* (B) in the leaves and roots of *P. stratiotes* exposed to heavy metals for 0, 1, and 4 h, normalized to β*-actin*.

## 4 Discussion

### 4.1 MT gene identification and evolutionary analysis

*P. stratiotes* is known for its ability to accumulate heavy metals from contaminated water, sequestering them primarily to its aerial parts (Lu *et al*., 2011; Zahari *et al*., 2021). This process is known to be associated with intracellular molecules such as MTs that play a critical role in metal detoxification (Gao *et al*., 2019; Talebi *et al*., 2019). While the accumulation of heavy metals by *P. stratiotes* is well demonstrated, the specific involvement of MTs in this process remains poorly understood. With the recent publication of the *P. stratiotes* chromosomes (Qian *et al*., 2022), a comprehensive genome-wide characterisation performed in this study successfully identified six novel MTs genes which were named *PsMT2f*, *PsMT3a-d*, and *PsMT4* according to Binz & Kägi (1999) and Cobbett & Goldsbrough (2002).

Using novel sequences as query sequences, high percentage identity and homology (**Table 3.1**-**3.2**) was observed with published MT sequences that are known to be involved in heavy metal tolerance and/or accumulation (Kim *et al*., 2011; Kim *et al*., 2012; Moyle *et al*., 2005). This high degree of homology often suggests functional similarities and has been widely used to characterise novel sequences, as demonstrated in several studies (Guo *et al*., 2008; Leszczyszyn *et al*., 2013; Pan *et al*., 2018; Parmar *et al*., 2012; Pearson, 2013; Yang *et al*., 2015). *PsMT2f* was mapped to chromosome 2, *PsMT3a* to chromosome 14, and *PsMT3b–d* were all located on chromosome 10 (**Figure 3.3**). The novel sequences contained a high proportion of predicted intronic regions that are known to often influence gene expression levels, enhance transcript stability, and protect coding regions from random mutations (Buchman & Berg, 1988; Das & Bansal, 2019; Jo & Choi, 2015). These intronic features may contribute to the functional diversity of MTs as observed in other plant species (Gao *et al*., 2022; Pan *et al*., 2018).

Furthermore, the high homology between novel MT3 sequences (*PsMT3a-d*) suggests the presence of multiple MT3 paralogs. Paralogs likely arose from gene duplication events in *P. stratiotes* (Qian *et al*., 2022). Duplication events are often essential for functional variation and adaptation in plants, providing them with the ability to survive under varying environmental conditions as observed in other plants (Ebadi *et al*., 2023; Gao *et al*., 2022; Pan *et al*., 2018; Qian *et al*., 2022; Zhou *et al*., 2019). The ratio of Ka/Ks ratio showed purifying selection (**Table 3.2**), essential for removing harmful mutations that often lead to nonsynonymous changes which renders proteins nonfunctional (Dong *et al*., 2019; Du *et al*., 2023; Roth & Liberles, 2006).

### 4.2 MT polypeptide sequence characteristics

The role of MTs in heavy metal chelation and ROS chelation relies on cysteine residues in the alpha and beta domains. These residues were preserved across all novel MTs (**Figure 3.1**), suggesting functional similarity with previously characterised MTs (Freisinger, 2011; Leszczyszyn *et al*., 2013; Ruttkay-Nedecky *et al*., 2013). Extra cysteine residues in the linker regions often improve heavy metal binding efficiency by providing extra ligands for heavy metal chelation (Parmar *et al*., 2012). Varying linker regions were observed, with some linker regions containing aromatic amino acids such as tyrosine, histidine, and phenylalanine, which can reduce the overall polypeptide charge by acting as a hydrogen bond donor or acceptor, and maintaining the high affinity of MTs for heavy metals (Blindauer, 2008; Leszczyszyn *et al*., 2013). This variation results in varying protein characteristics, as the pI of novel MTs were in the range of 4.49-6.94, and hydropathy index in the range of - 0.41-0.08 (**Table 3.1**). These variations in pI and hydropathy index are often associated with differences in protein function, as they influence how MTs respond to factors such as pH and solubility under varying physiological conditions (Di Rienzo *et al*., 2021; Tokmakov *et al*., 2021). Furthermore, using DeepLoc, novel MTs were shown to be located in different compartments. *PsMT2f* was predicted to be located in cytoplasm while *PsMT3a-d* and *PsMT4* were predicted to be extracellular proteins (Goldberg *et al*., 2014), similar to what was observed in other plants such as *Nicotiana Tabacum* (Ren *et al*., 2012; Yu *et al*., 2021). MTs localization within different cellular compartments can affect how MTs interact with various cellular processes (Parameswari *et al*., 2021; Vašák & Meloni, 2011).

Using MEME online tool, MT3 sequences had conserved motifs 1, 2, and 3 while *PsMT2f* and *PsMT4* had several motifs (**Figure 3.4**). The results presented here show that while certain motifs are conserved and suggest a shared functional role, there is also predicted functional diversity among MTs due to motif variations necessary to cater to the specific needs of organisms (Hassinen *et al*., 2011). This functional diversity is crucial for the adaptation and specialisation of different MTs in various biological contexts of *P. stratiotes*, with motif variation potentially reflecting differences in metal-binding properties and stress responses. Such information may be valuable for determining whether specific MTs contribute to enhanced tolerance or accumulation of particular contaminant metals, thereby informing the practical application of *P. stratiotes* in phytoremediation.

### 4.3 Kozak sequences

The Kozak sequence plays a crucial role in translation initiation as it enhances the binding of the 40S ribosomal subunit to the start codon, ensuring the accurate assembly of the ribosome (Hernández *et al*., 2019; Kim *et al*., 2014; Kozak, 1986). Bases at -3 (R) and +4 (G) from the nucleotide base adenine (A) of the TSS were observed to be universally conserved among most eukaryotic genes, suggesting that they play an important role in initiating translation (Hernández *et al*., 2019). Analysis of Kozak sequences showed that *PsMT2f* and *PsMT3a-d* had conserved –3 (R) and +4 (T), and *PsMT4* had conserved –3 (R) and +4 (G) (**Table 3.1**). This is not uncommon as it has been observed in MTs from various plants (Ambrosini *et al*., 2022; Hernández *et al*., 2019; Kaur *et al*., 2020; Kozak, 1987; Kozak, 1986; Meijer & Thomas, 2002). The presence of +4 (T) instead of +4 (G) often suggest reduced translation efficiency, which can be a regulatory effect on the translation of MTs (Acevedo *et al*., 2018; Kaur *et al*., 2020; Kozak, 1987).

### 4.4 Real time PCR optimization and reference gene analysis

The response of plants to heavy metal stress involves changes in the transcriptional and translational mechanisms that result in the production of several molecules, including metal-chelating biomolecules (Hassinen *et al*., 2010; Lv *et al*., 2012; Saeed *et al*., 2020). The novel primers designed for relative gene expression analysis successfully amplified the target genes, as demonstrated in **Figure 3.5**. The amplification efficiencies fell within the expected range of 100 ± 10%, aligning with established guidelines (Ginzinger, 2002; Livak & Schmittgen, 2001). These results indicate that the novel primers exhibited high specificity, sensitivity, and efficiency, which are key parameters required for reliable and accurate gene expression studies.

For relative quantification by real-time PCR, multiple reference genes are required for gene expression normalization. Reference genes should be constitutively expressed and non-regulated, and their expression should be independent of the environmental conditions (Chen *et al*., 2021; Sen *et al*., 2021). Reference genes are usually metabolic genes involved in basal cellular activities such as cellular structure maintenance, protein translation, and carbon metabolism (Samarth & Jameson, 2019). Several studies have used endogenous *18S rRNA*, *EF 1*α, or β*-actin* genes as reference genes in plants exposed to heavy metals (Hong *et al*., 2008; Karimi & Mohsenzadeh, 2017; Wang *et al*., 2017). Some studies have shown poor expression stability of candidate reference genes such as β*-actin* under abiotic stress (Gu *et al*., 2011; Zhao *et al*., 2020). This shows that the expression and stability of reference genes vary between plants and plant tissues; hence, there is a need to investigate the stability of reference genes each time a plant is exposed to different experimental conditions (Song *et al*., 2020; Wan *et al*., 2017; Yi *et al*., 2012).

To date, research regarding reference genes for *P. stratiotes* has been quite limited. In this study, of the four candidate reference genes that were selected, and only two (*18S rRNA* and β*-actin*) were constitutively expressed in *P. stratiotes* at sufficient levels (**Table 3.4**). Although *18S rRNA* has been widely used as a reference gene in plants exposed to various abiotic stresses caused by heavy metals (Zhao *et al*., 2020; Chen *et al*., 2021; Sen *et al*., 2021), a high SD of 3.28 in leaves and 4.37 in roots, along with a CV of 33.32% and 38.77% (**Figure 3.6**) have been observed, suggesting the instability of *18S rRNA* expression in *P. stratiotes*. This is not uncommon, as several studies have also noted the limitations of using *18S rRNA* as the reference gene (Bisht *et al*., 2021; Ebrahimi *et al*., 2024; Kim *et al*., 2010; Yi *et al*., 2012).

In our analysis using geNorm (**Table 3.5**), the obtained M value was >1.5 with both genes, indicating that at least one of the genes was unstable under experimental conditions (Vandesompele *et al*., 2002). BestKeeper showed an SD of >1. However, β*-actin* had the lowest SD in both leaves and roots when compared to 18S rRNA, suggesting that it had better stability than 18S rRNA (Köhsler *et al*., 2020; Pfaffl *et al*., 2004). Using NormFinder, the stability value obtained between β*-actin* and *18S rRNA* was 1.02 and 1.86 in roots and leaves, respectively, and the comprehensive ranking method, which combines the results from all three statistical tools showed that β*-actin* was the most stable gene in *P. stratiotes*. Based on this consistent performance across multiple statistical tools, β*-actin* was selected as the reference gene for all subsequent gene expression analyses (Andersen *et al*., 2004; Linardić & Braybrook, 2021).

### 4.5 Expression profile of MTs under heavy metal stress

*PsMT2f* and *PsMT3b-d* were shown to be constitutively expressed in *P. stratiotes,* suggesting that they play a prominent role in plant growth and development and in maintaining homeostasis. This observation aligns with previous studies that have highlighted the involvement of MTs in metabolism regulation of various plants (Joshi *et al*., 2016; Leszczyszyn *et al*., 2013). In addition, the constitutive expression of *PsMT2f* and *PsMT3b-d* indicates their preparedness to respond to environmental stresses that frequently lead to the production of ROS that can cause significant damage to cellular structures, including proteins, lipids, and DNA (Guo *et al*., 2023; Mansoor *et al*., 2022).

At 1 h, *PsMT2f* expression remained unchanged in both roots and leaves, whereas *PsMT3b-d* showed no change in roots but was upregulated in leaves (**Figure 3.7**). No changes in the expression of *PsMT2f* may be attributed to the fact that they are constitutively expressed more than 10-fold greater than *PsMT3b-d*, therefore, already-made MTs may be available to bind and sequester Cu within the first hour without the need for upregulating *PsMT2f* expression. Variation in MTs expression can also be attributed to the differences in the regulatory mechanisms controlling these genes, allowing *PsMT3b-d* to be transcriptionally activated while *PsMT2f* remains unchanged.

Differences in the expression rates of MTs from the same plant are not uncommon, as previously documented. In *Brassica rapa*, *BrMT1* and *BrMT2* were not induced, whereas *BrMT3* expression increased by approximately 2-fold after exposure to Fe. Furthermore, in the presence of Cu, *BrMT2* was downregulated, *BrMT3* expression remained constant, and *BrMT1* was upregulated (Ahn *et al*., 2012). The expression of MTs, such as *PsMT2f* and *PsMT3b-d*, depends on their physiological roles that are predicted to include heavy metal chelation, regulation of ion homeostasis, and quenching of ROS (Cobbett & Goldsbrough, 2002; Joshi *et al*., 2016). These roles depend on various factors that are not fully understood. However, DeepLoc analysis showed that *PsMT2f* is likely to be an intracellular protein, whereas *PsMT3b-d* are proposed extracellular proteins. As a first step towards dealing with metal intoxication, plants often adopt an avoidance strategy to restrict the accumulation of heavy metals through several mechanisms, including immobilization and complexation by root exudates. It is reasonable to suggest that extracellular proteins may be expressed earlier than intracellular proteins, as their role in binding heavy metals could help mitigate absorption into plant cells, and this needs to be further validated through temporal expression and localisation studies.

Following that, at 4 h, *PsMT2f* was downregulated in the roots and upregulated in the leaves, while *PsMT3b-d* expression remained unchanged in both tissues (**Figure 3.7**). Heavy metal exposure in the roots has been shown to induce metal stress, resulting in growth impairment and cell death (Yadav *et al*., 2021). Heavy metal stress also causes root loss in *P. stratiotes* (Farnese *et al*. 2014), therefore, the downregulation and marginal decrease in expression observed at 4 h timepoint can be attributed to changes in the physiological state of the roots due to heavy metal exposure. This is not uncommon, as downregulation of MTs has been observed in other plants (Duan *et al*., 2019; Liu *et al*., 2021). An increase in the expression of *PsMT2f* observed in the leaves is likely essential for the production of polypeptides that help curb heavy metal stress by binding and sequestering them in the aerial parts of the plant (Alsafran *et al*., 2022; Sharma *et al*., 2016). Similar results have been observed with MTs from various plants, including *A. thaliana* (Guo *et al*., 2003), *S. officinarum* (Guo *et al*., 2013), *S. lycopersicum* (Bolukbasi, 2021), and *M. domestica* (Wan *et al*., 2019).

As *PsMT2f* polypeptide contains cysteine residues that are known for binding and chelating heavy metals, the observed increase in *PsMT2f* gene expression may be indicative of an adaptive response in *P. stratiotes* to heavy metal stress (Joshi *et al*., 2016). This suggests that *PsMT2f* could be part of a defense mechanism, likely facilitating not only metal sequestration but also helps maintain cellular homeostasis, enhancing the plant’s overall tolerance to heavy metal stress (Joshi *et al*., 2016; Leszczyszyn *et al*., 2013). Its expression at 4 h in the leaves can be linked to its proposed intracellular function. After 4 h of exposure, studies have shown that plants would have accumulated and translocated heavy metals to different plant tissues (Jamil *et al*., 1985; Kajala *et al*., 2019); hence, there is a need for intracellular mechanisms to bind and chelate heavy metals. Since *PsMT2f* is a likely cytoplasmic protein, similar to *NtMT2a*, *NtMT2b*, *NtMT2d*, and *NtMT2f-g* from *Nicotiana Tabacum* (Yu *et al*., 2021), upregulation of *PsMT2f* at the 4 h time point might be essential to produce *PsMT2f* polypeptides for heavy metal chelation.

Overall, the responses of *PsMT2f* and *PsMT3b–d* to heavy metal exposure demonstrated distinct expression patterns, suggesting gene-specific roles in the heavy metal stress response of *P. stratiotes*. These differences in expression may stem from various underlying factors, including differences in gene regulation, the presence of specific cis-regulatory elements, and unique sequence characteristics inherent to each gene. Additionally, the differential expression profiles reflect the complexity of the plant’s adaptive biochemical and molecular mechanisms to cope with metal-induced stress. This study reinforces the involvement of MT genes in heavy metal tolerance and provides a foundation for future functional analyses aimed at understanding their precise roles in phytoremediation and stress mitigation in *P. stratiotes*.

### 5 Conclusion

In the present study, genome-wide analysis was used to identify six novel putative MTs in *P. stratiotes*. One was classified as type two (*PsMT2f*), four were paralogs and were classified as type three (*PsMT3a-d*), and one was classified as type four (*PsMT4*). The analyses performed showed a close association between novel MTs with MTs from various plants. Novel sequences have regulatory elements involved in plant development, hormonal response, light response, and stress response, and Kozak sequence analysis revealed the presence of adequate and strong TSS, essential for transcription initiation. Novel primers were designed and optimized, and β*-actin* reference gene was shown to be the most stable reference gene for use in *P. stratiotes* exposed to Cu. Relative gene expression by real time PCR showed a distinct response pattern of stress related genes, *PsMT2f* and *PsMT3b-d*, in *P. stratiotes* exposed to Cu, and confirms their response to heavy metal related stress in *P. stratiotes*. Examining the responses of novel MTs to heavy metal exposure improves our understanding of biomolecular mechanisms that facilitate metal tolerance and accumulation *P. stratiotes*.

